# Scalable Multi-Sample Single-Cell Data Analysis by Partition-Assisted Clustering and Multiple Alignments of Networks

**DOI:** 10.1101/116566

**Authors:** Ye Henry Li, Dangna Li, Nikolay Samusik, Xiaowei Wang, Leying Guan, Garry P. Nolan, Wing Hung Wong

## Abstract

Mass cytometry (CyTOF) has greatly expanded the capability of cytometry. It is now easy to generate multiple CyTOF samples in a single study, with each sample containing single-cell measurement on 50 markers for more than hundreds of thousands of cells. Current methods do not adequately address the issues concerning combining multiple samples for subpopulation discovery, and these issues can be quickly and dramatically amplified with increasing number of samples. To overcome this limitation, we developed Partition-Assisted Clustering and Multiple Alignments of Networks (PAC-MAN) for the fast automatic identification of cell populations in CyTOF data closely matching that of expert manual-discovery, and for alignments between subpopulations across samples to define dataset-level cellular states. PAC-MAN is computationally efficient, allowing the management of very large CyTOF datasets, which are increasingly common in clinical studies and cancer studies that monitor various tissue samples for each subject.

**Author Summary:** Recently, the cytometry field has experienced rapid advancement in the development of mass cytometry (CyTOF). CyTOF enables a significant increase in the ability to monitor 50 or more cellular markers for millions of cells at the single-cell level. Initial studies with CyTOF focused on few samples, in which expert manual discovery of cell types were acceptable. As the technology matures, it is now feasible to collect more samples, which enables systematic studies of cell types across multiple samples. However, the statistical and computational issues surrounding multi-sample analysis have not been previously examined in detail. Furthermore, it was not clear how the data analysis could be scaled for hundreds of samples, such as those in clinical studies. In this work, we present a scalable analysis pipeline that is grounded in strong statistical foundation. Partition-Assisted Clustering (PAC) offers fast and accurate clustering and Multiple Alignments of Networks (MAN) utilizes network structures learned from each homogeneous cluster to organize the data into data-set level clusters. PAC-MAN thus enables the analysis of a large CyTOF dataset that was previously too large to be analyzed systematically; this pipeline can be extended to the analysis of similarly large or larger datasets.

## Introduction

Analyses of CyTOF data rely on many of the tools and ideas from flow cytometry (FC) data analysis, as CyTOF datasets are essentially higher dimensional versions of flow cytometry datasets. Currently, the most widely used method in FC is still human hand-gating, as other methods often fail to extract meaningful subpopulations of cells automatically. In hand-gating, we draw polygons or other enclosures around pockets of cell events on a two-dimensional scatterplot to define subpopulations and cellular states that are observed in the data. This process is painfully time-consuming and requires advance knowledge of the marker panel design, the quality of the staining reagents, and, most importantly, *a priori* what cell subpopulations to expect to occur in the data. When presented with a new set of marker panels and biological system, the researcher would find it difficult to delineate the cell events, especially in high-dimensional and multi-sample datasets.

The inefficient nature of hand-gating in flow cytometry motivated algorithmic development in automatic gating. Perhaps the most popular is flowMeans[1], which is optimized for FC and can learn subpopulations in FC data[2] in an automated manner; however, it has not been successfully applied to CyTOF data analysis. Currently, most data analysis tools created for flow cytometry data analyses are not easily applicable for high-dimensional datasets[3]. An exception is SPADE, which was developed and optimized specifically for the analysis of CyTOF datasets[3]. flowMeans and SPADE constitute the leading computational methods in cytometry, but as shown later in this work, their performance may become sub-optimal when challenged with large and high-dimensional datasets. There are also other recent clustering-based tools that utilize dimensionality reduction and projections of high-dimensional data, however, these tools do not directly learn the subpopulations for all the cell events, and may be too slow to complete data analysis for an increasing amount of samples.

In this study, we address the data analysis challenges in two major steps. First, we propose the partition-assisted clustering (PAC) approach, which produces a partition of the k-dimensional space (k=number of markers) that captures the essential characteristic of the data distribution. This partitioning methodology is grounded in a strong mathematical framework of partition-based high-dimensional density estimation[4–8]. The mathematical framework offers the guarantee that these partitions approximate the underlying empirical data distribution; this step is faster than the recent k-nearest neighbor-based method [9] and is essential to the scalability of our clustering approach to analyze datasets with many samples. The clustering of cells based on recursive partitioning is then refined by a small number of k-means style iterations before a merging step to produce the final clustering.

Secondly, the subpopulations learned separately in multiple different but related datasets can be aligned by marker network structures (multiple alignments of networks, or MAN), making it possible to characterize the relationships of subpopulations across different samples automatically. The ability to do so is critical for monitoring changes in a subpopulation across different conditions. Importantly, in every study, batch effect is present; batch effects shift subpopulation signals so that the means can be different from experiment to experiment. PAC-MAN naturally addresses batch effects in finding the alignments of the same or closely related subpopulations from different samples.

PAC-MAN finds homogeneous clusters efficiently with all data points in a scalable fashion and enables the matching of these clusters across different samples to discover cluster relationships in the form of clades.

## Results and Discussion

### PAC

PAC has two parts: partitioning and post-processing. In the partitioning part of PAC, the data space is recursively divided into smaller hyper-rectangles based on the number of data points in the locality (Fig 1a). The partitioning is accomplished by either Bayesian Sequential Partition (BSP) with limited look-ahead (Fig 1a and 1b) or Discrepancy Sequential Partition (DSP) (Fig 1a); these are two fast variants of partition-based density estimation methods previously developed by our group [4–8], with DSP being the fastest. BSP and DSP divide the sample space into hyper-rectangles with uniform density value in each of them. The subsetting of cells according to the partitioning provides a principled way of clustering the cells that reflects the characteristics of the underlying distribution. In particular, each significant mode is captured by a number of closely located rectangles with high-density values (Fig 1c). Although this method allows a fast and unbiased localization of the high-density regions of the data space, we should not use the hyper-rectangles directly to define the final cluster boundaries for two reasons. First, real clusters are likely to be shaped elliptically, therefore, the data points in the corners of a hyper-rectangle are likely to be incorrectly clustered. Second, a real cluster is often split into more than one closely located high-density rectangles. We designed post-processing steps to overcome these limitations: 1) a small number of k-means iterations is used to round out the corners of the hyper-rectangles, 2) a merging process is implemented to ameliorate the splitting problem, which is inspired by the flowMeans algorithm. The details of post-processing are given in the Materials and Methods. The resulting method is named b-PAC or d-PAC depending on whether the partition is produced by BSP or DSP.

**Fig 1:**
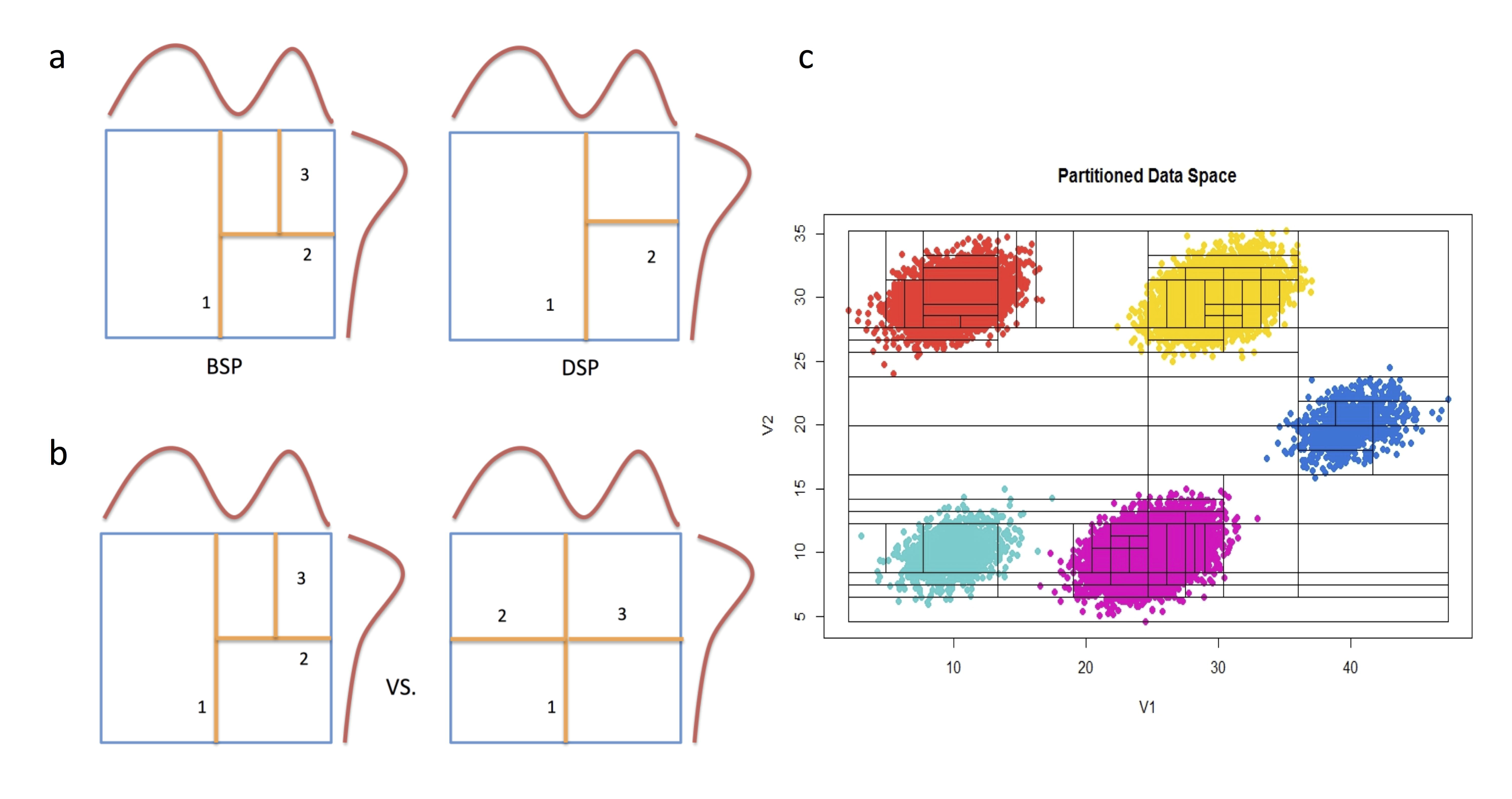
PAC recursively partitions the data space to obtain rational initialization structure. (a) Partition-based methods estimate data density by cutting the data space into smaller rectangles. Bayesian Sequential Partition (BSP) divides the data space via binary partition in the middle of the bounded region, while that of Discrepancy Sequential Partition (DSP) occur at the location that balances the data point uniformly on both sides of the cut. The numbers denote sequential order of partitions. Since DSP adapts to the data points, it converges on the estimated density faster than BSP. (b) In the (one-step) look-ahead of version of partition, the algorithm cuts the data space for all potential cuts plus one step more (steps 2 and 3), and it finds the optimal future version (after step 3), which determines the actual cut (step 2). (c) The partitioning of simulated data space containing five subpopulations; the hyper-rectangles surround high-density areas, approximating the underlying distribution.

### MAN

An approach to analyze multiple related samples of CyTOF data is to pool all samples into a combined sample before detection of subpopulations. This is a natural approach under the assumptions that there are no significant batch effects or systematic shifts in cell subpopulations across the different samples. However, such assumptions may not hold due to one or more of the following reasons:

1. *Dataset size and instruments used*. Large number of samples usually means the samples were collected on different days with different experimental preparations. Many steps can introduce significant shifts in measurement levels.
2. *Staining reagents*. Reagents such as antibodies, purchased from different vendors and batch preparations can affect the overall signal. While saturation of reagents in the protocol could help eliminate the batch effects in the staining procedure, this approach is costly and might not work for all antibodies, especially those with poor specificity.
3. *Normalization beads stock*. While normalization beads[10] help to control for the signal level, especially within one experiment, the age of the beads stock and their preparation could lead to significant batch effects. In addition, there are different types of normalization beads and normalization calculations.
4. *Human work variation*. While many researchers are studying the same system (e.g., immune system), different protocols and implementation by different researchers, who sometimes perform experimental steps slightly differently, can lead to batch effects.
5. *Subpopulation dynamics*. The subpopulation centers can move from sample to sample due to treatments on the cells in treatment-control studies or perturbation studies. General practice is to cluster by phenotypic markers.
6. *Sample background*. If the data came from different cell lines or individuals in a clinical study, the measurement levels and proportions of cell subpopulations would be expected to change from sample to sample. Without expert scrutiny, it would be difficult to make sense of the data with current data analysis tools.

Could we extract shared information that allows us to interpret cross-sample similarities and differences? To ameliorate these difficulties, we have designed an alternative approach that is effective in the presence of substantial systematic between-sample variation. In this approach, each sample is analyzed separately (by PAC) to discover within-sample subpopulations. Over-partitioning in this step is allowed in order not to miss small subpopulations in high dimension due to lack of prior knowledge. The subpopulations from all samples are then compared to each other based on a pairwise dissimilarity measure designed to capture the differences in within-sample distributions (among the markers) across two subpopulations. Using this dissimilarity, we perform bottom-up hierarchical clustering of the subpopulations to represent the relationship among the subpopulations. The resulting tree of subpopulations is then used to guide the merging of subpopulations from the same sample, and to establish linkage of related subpopulations from different samples. We note that the design of a dissimilarity measure (Materials and Methods) that is not sensitive to systematic sample-to-sample variation is a novel aspect of our approach. The merging of subpopulations from the same sample is also important, as it offers a way to consolidate any over-partitioning that may have occurred during the initial PAC analysis of each sample. We emphasize that, as with the usage of all statistical methods, the user must utilize samples or datasets that are considered as good as possible; interpretation of the analysis results rely on the researchers to collect data with validated reagents for all samples.

### Rational initialization for PAC increases clustering effectiveness

Appropriate initialization of clustering is very important for eventually finding the optimal clustering labels; PAC works well because the implicit density estimation procedure yields rational centers to learn the modes of sample subpopulations. When tested on the hand-gated CyTOF data on the bone marrow sample in (14), compared to k-means alone, PAC gives lower total sums of squares and higher F-measures in the subpopulations (Fig 2a and 2b). This process also helps PAC to converge in 50 iterations (Fig 3) in post-processing, whereas k-means performs very poorly even after 5000 iterations (Fig 4). Through the lens of t-sne plots (Fig 4), the PAC results are more similar to the hand-gating results, while the k-means, flowMeans, and SPADE clustering results perform poorly. In flowMeans, several large subpopulations are merged. SPADE’s separation of points is inconsistent and highly heterogeneous, probably due to its down-sampling nature. On the other hand, by inspection, PAC obtains similar separation for both the major and minor subpopulations as the hand-gating results.

**Fig 2.**
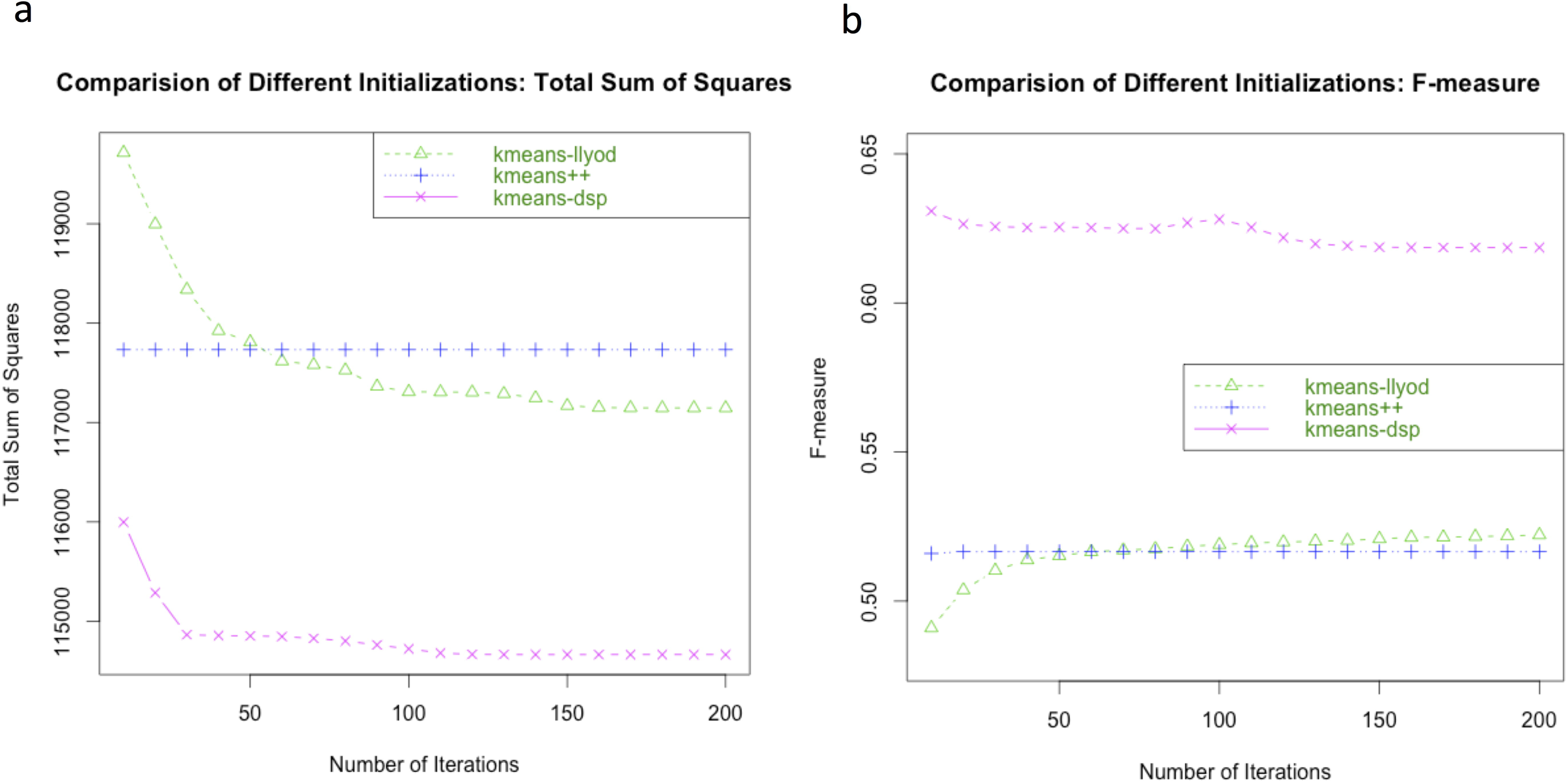
Rational initialization is better than random initialization. The hand-gated CyTOF data (see S1 Fig) is used for illustration. In this case, (a) the overall sum of squares error is lower and (b) the F-measure is higher for PAC.

**Fig 3.**
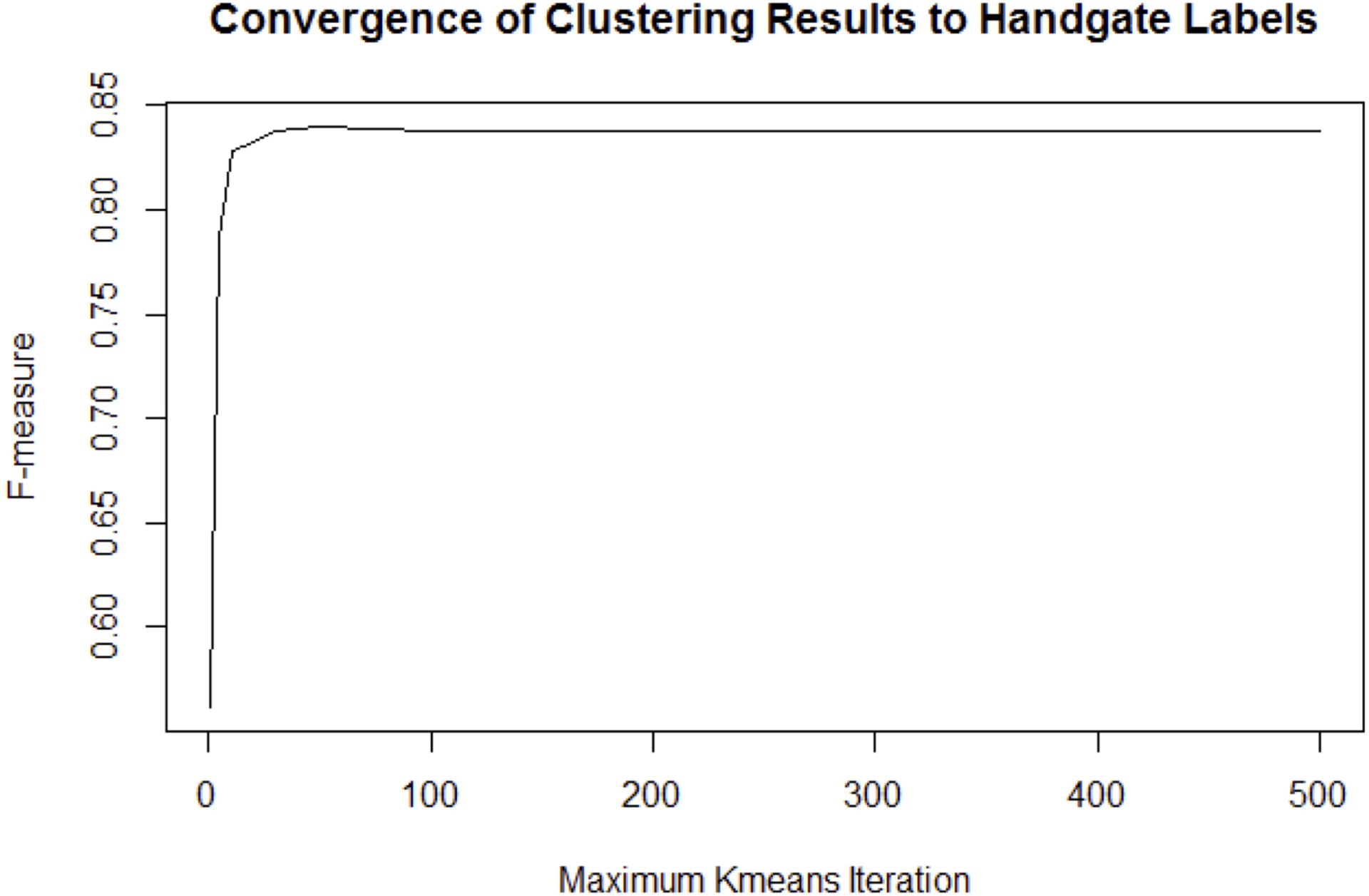
Rational initialization and minimal kmeans post-processing iterations give fast convergence. The convergence of PAC toward the hand-gated results, or ground truth, is fast. It takes less than 50 downstream post-processing kmeans iterations for the PAC to achieve convergence.

**Fig 4.**
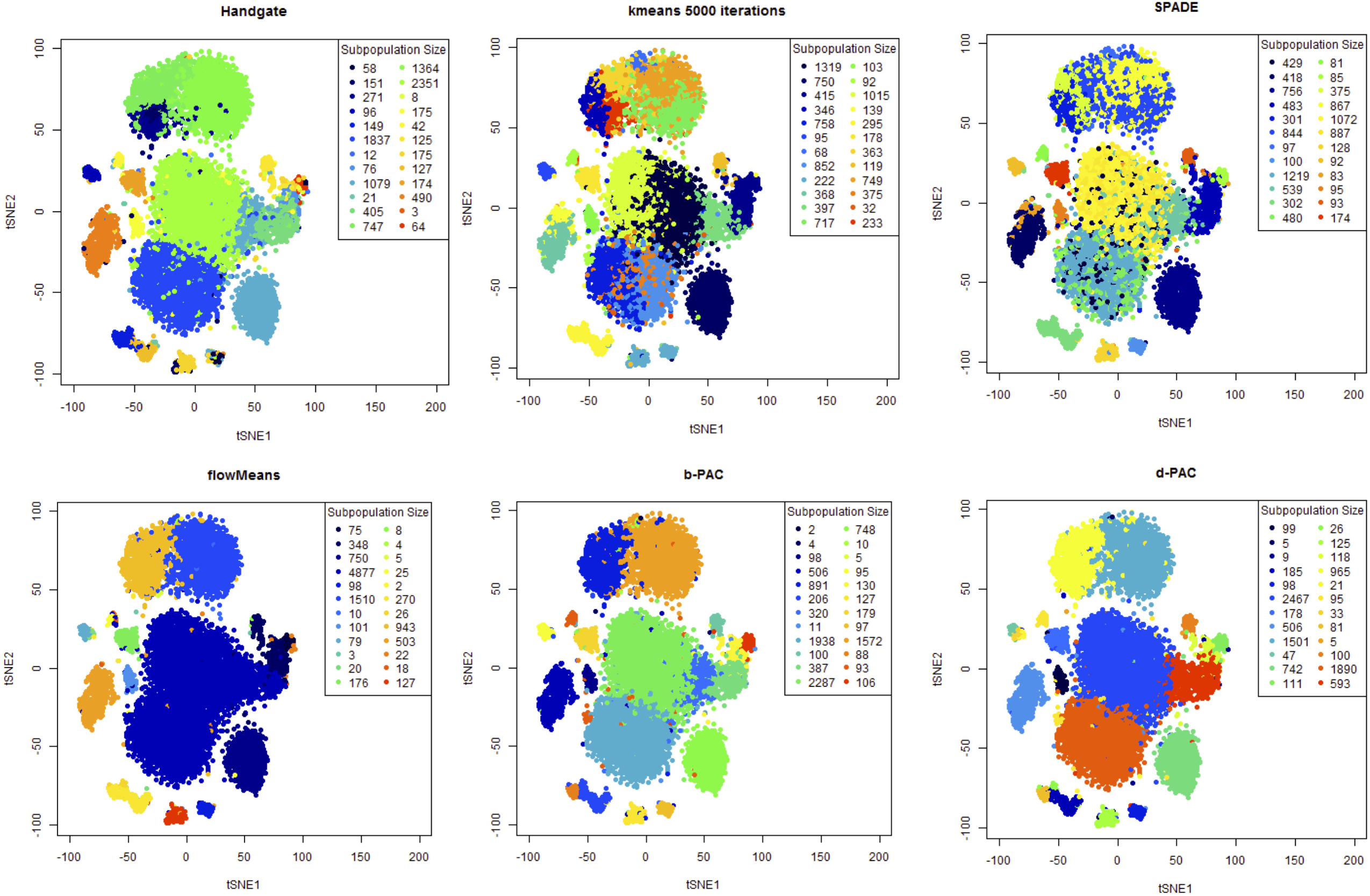
Visualization and comparison of clustering results by t-sne plots. Each t-sne plot contains the same 10,000 cell events from the hand-gated CyTOF data with different set of colored labels drawn. Note that the colors are informative only within each panel. These labels are from kmeans, SPADE, flowMeans, b-PAC, and d-PAC. The subpopulation numbers for all methods were set to be the same as that of hand-gated results. PAC methods achieve a significantly better convergence to the hand-gate labels than alternative methods.

### PAC is consistently better than flowMeans and SPADE for simulated datasets and hand-gated cytometry datasets

In the systematic simulation study, we challenged the methods with different datasets with varying number of dimensions, number of subpopulations, and separation between the subpopulations. The F-measure and p-measures for the PAC methods are consistently equal or higher than that of flowMeans and SPADE (Table 1 and S2a Fig). In addition, we observe that flowMeans gives inconsistent F-measures for similar datasets (Table 1), which may be due to the convergence of k-means to a local minimum without a rational initialization.

**Table 1.**
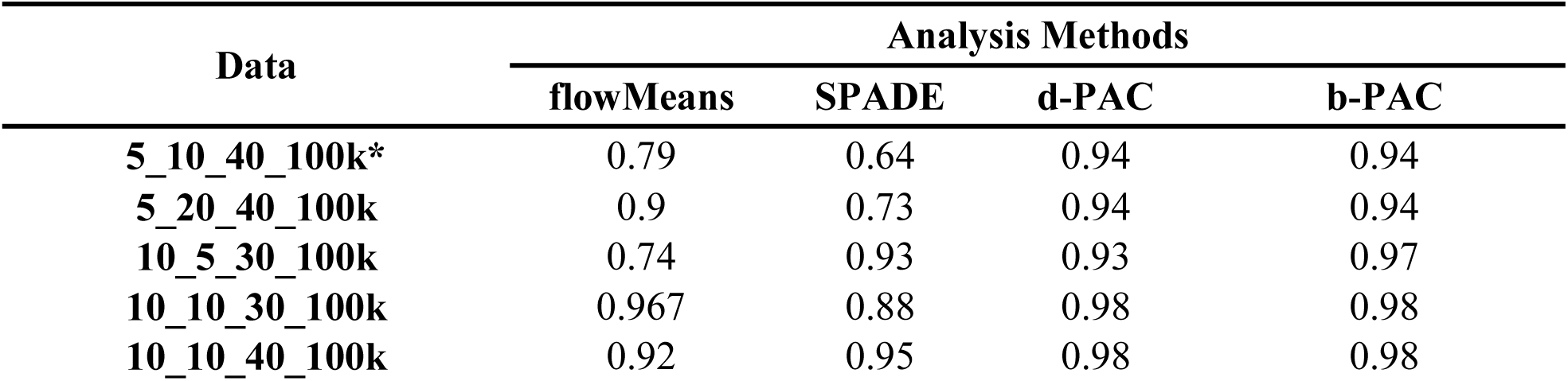
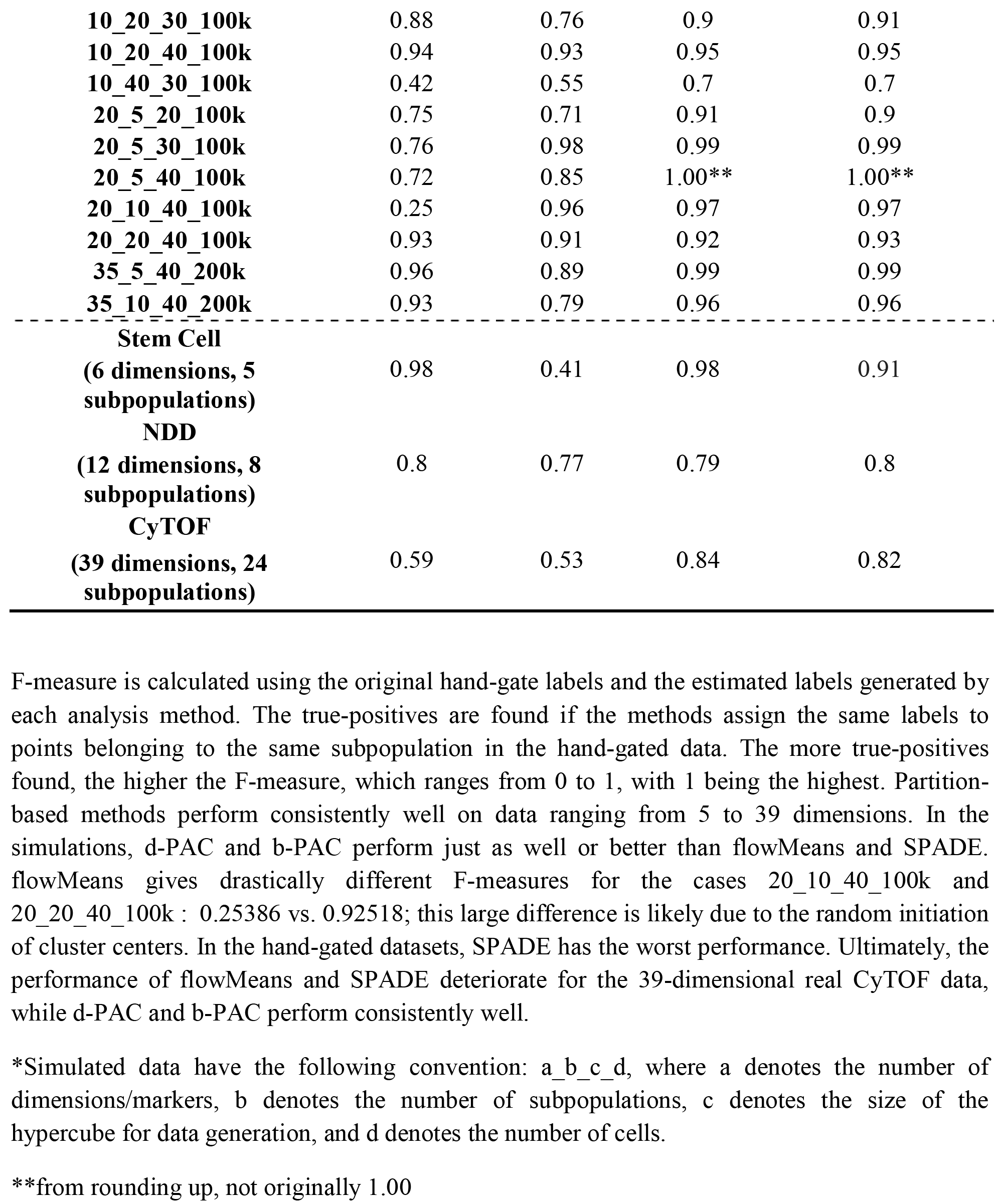
F-measure Comparisons of Methods on Simulated and Hand-gated Cytometry Datasets.

Next, we tested the methods based on published hand-gated cytometry datasets to see how similar the estimated subpopulations are to those obtained by human experts. We applied the methods on the hematopoietic stem cell transplant and Normal Donors datasets from the FlowCAP challenges[2] and on the subset of gated mouse bone marrow CyTOF dataset (Dataset 5) recently published[11]. The gating strategy of the CyTOF dataset is provided in Fig S1. The dataset and expert gating strategy are the same as described earlier[12]. Note that in the flow cytometry data, the computed F-measures are slightly lower than that reported in FlowCAP; this is due to the difference in the definition of F-measures. Overall, the PAC outperforms flowMeans and SPADE by consistently obtaining higher F-measures (Table 1). In particular, in the CyTOF data example, PAC generated significantly higher F-measures (greater than 0.82) than flowMeans and SPADE (0.59 and 0.53, respectively). In addition, PAC gives higher overall subpopulation-specific purities (S2b Fig and S1 Table). These results indicate that PAC gives consistently good results for both low and high-dimensional datasets. Furthermore, PAC results match human hand-gating results very well. The consistency between PAC-MAN results and hand-gating results in this large data set confirms the practical utility of the methodology.

### Separate-then-combine outperforms pool approach when batch effect is present

It is natural to analyze samples separately then combine the subpopulation features for downstream analysis in the multiple samples setting. However, we need to resolve the batch effects. Two distinct subpopulations could overlap in the combined/pooled sample, such as in the case when the data came from two generations of CyTOF instruments (newer instrument elevates the signals). On the other hand, in cases with changing means, two subpopulations can evolve together such that their means change slightly, but enough to shadow each other when samples are merged prior to clustering.

We introduce Multiple Alignments of Networks to resolve the management issue surrounding the organization of homogeneous clusters found in the PAC step (Fig 5). First, we consider the overlapping scenario (Fig 6a). When viewed together in the merged sample, the right subpopulation from sample 1 overlaps with the left subpopulation in sample 2 (Fig 6b left panel). There is no way to use expression level alone to delineate the two overlapping subpopulations (Fig 6b right panel). By learning more subpopulations using PAC, there are some hints that multiple subpopulations are present (Fig 6c). Despite these hints, it would not be possible to say whether the shadowed subpopulations relate in any way to other distinct subpopulations.

**Fig 5.**
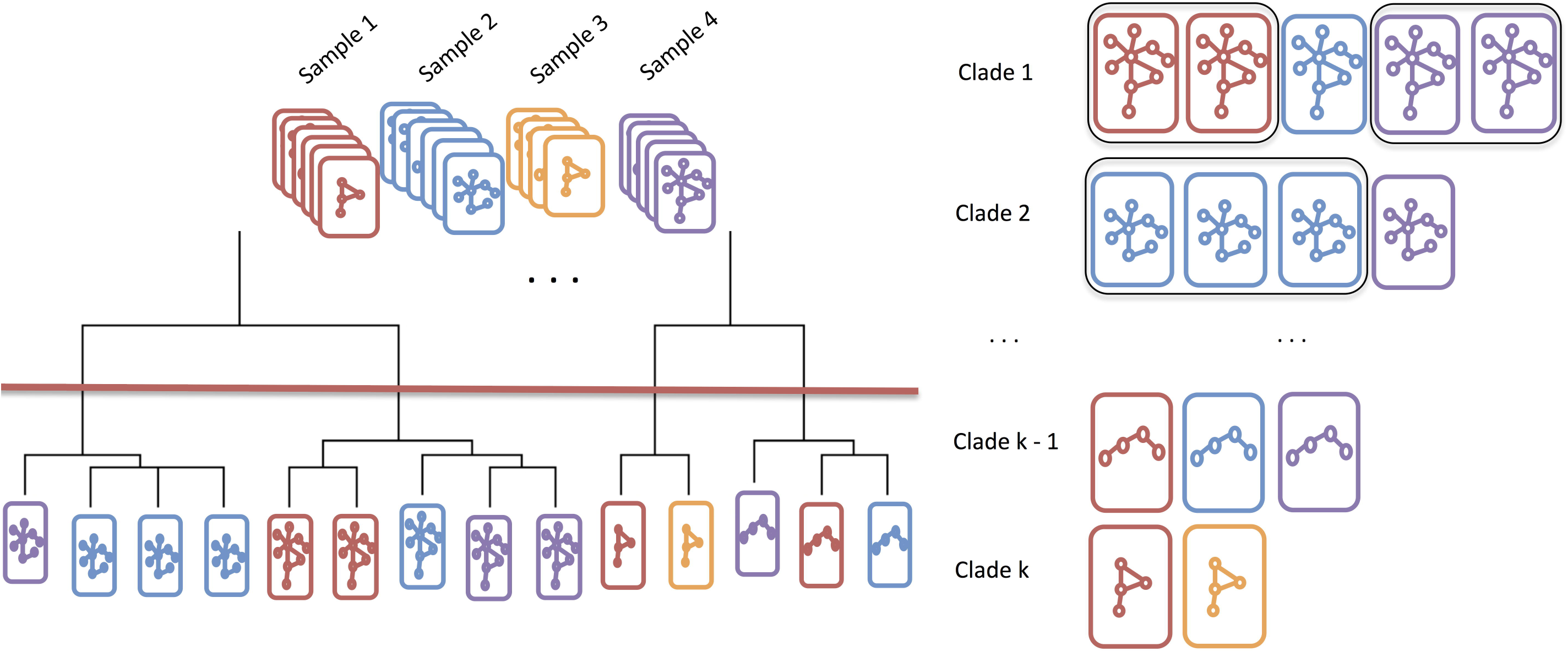
Schematic analogy of MAN. Consider a deck of networks (in analogy to cards), with each “suit” representing a sample and each “rank” representing a unique network structure. The networks are aligned by similarity and organized on a dendrogram. The tree is cut (red line) at the user-specified level to output the desired *k* clades. Within each clade, the network structures are similar or the same. If the same sample has multiple networks in the same clade, then these networks are merged (black box around same cards).

**Fig 6.**
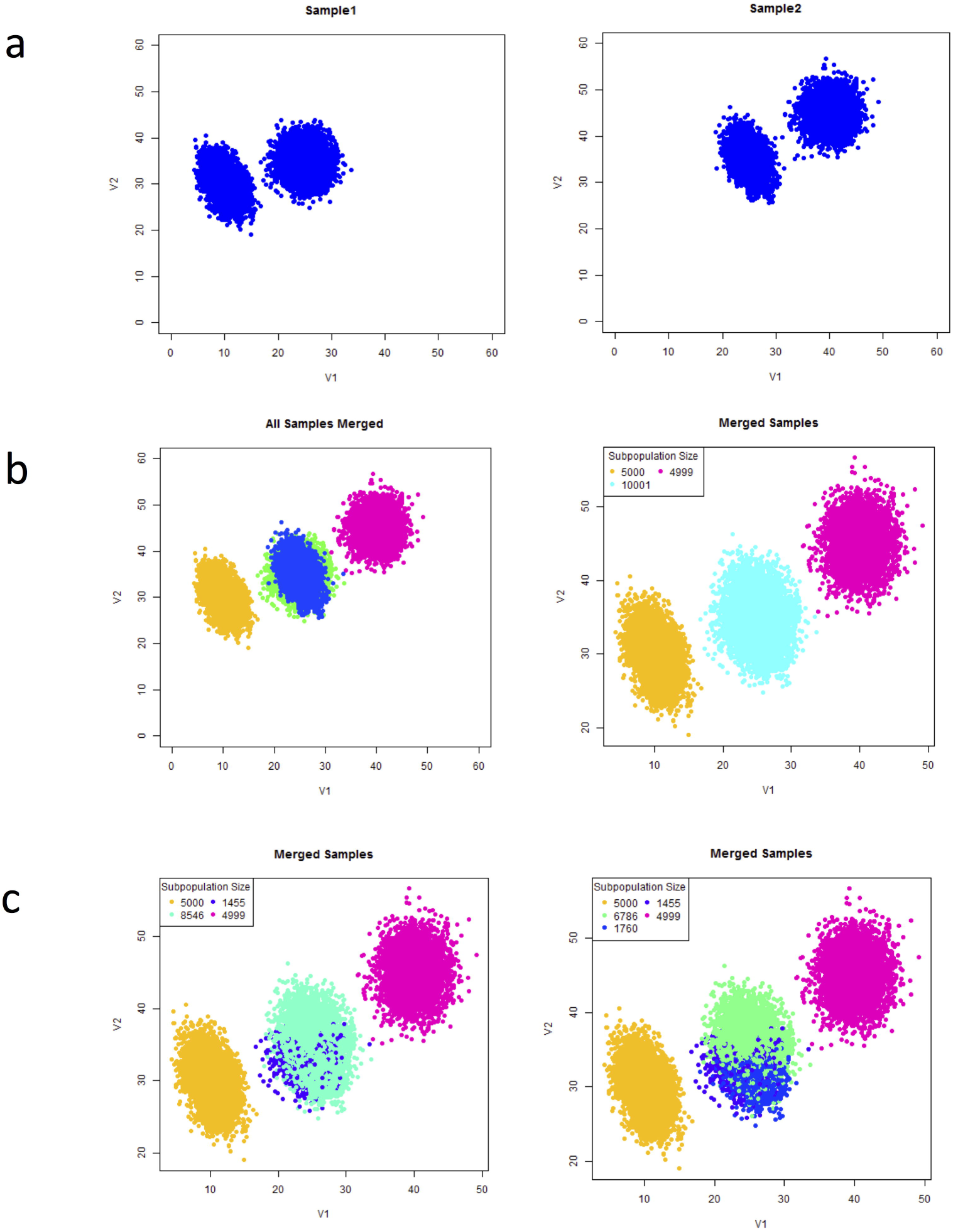
Simple Batch Effect Scenario. (a) Simulated data samples with two of the same subpopulations. The means shifted due to measurement batch effect. (b) When the samples are combined, as in the case of analyzing all samples together, two different subpopulations overlap (left panel). The overlapped subpopulations cannot be distinguished by clustering (right panel). (c) PAC could be used to discover more subpopulations, however, the hints of the present of another subpopulation do not help to resolve the batch effect.

PAC-MAN resolves the overlapping issue by analyzing the samples separately (Fig 7). In the case in which we do not know *a priori* the number of true subpopulations, we learn three subpopulations per sample (Fig 7a). The network structures of the subpopulations discovered are presented in Fig 7b-c and we see that the third subpopulations from the two samples share the same network structures, while the first subpopulations of the two samples differ by only one edge; these respective networks are clustered together in the dendrogram (Fig 8a right panel). By utilizing the networks, the clades that represent the same and/or similar subpopulations of cells can be established. Clustering by network structures alone resolves the majority of points in the data (Fig 8a, left panel). Furthermore, as discussed next, by incorporating marker levels into the alignment process, all the subpopulations can be resolved (Fig 8b).

**Fig. 7.**
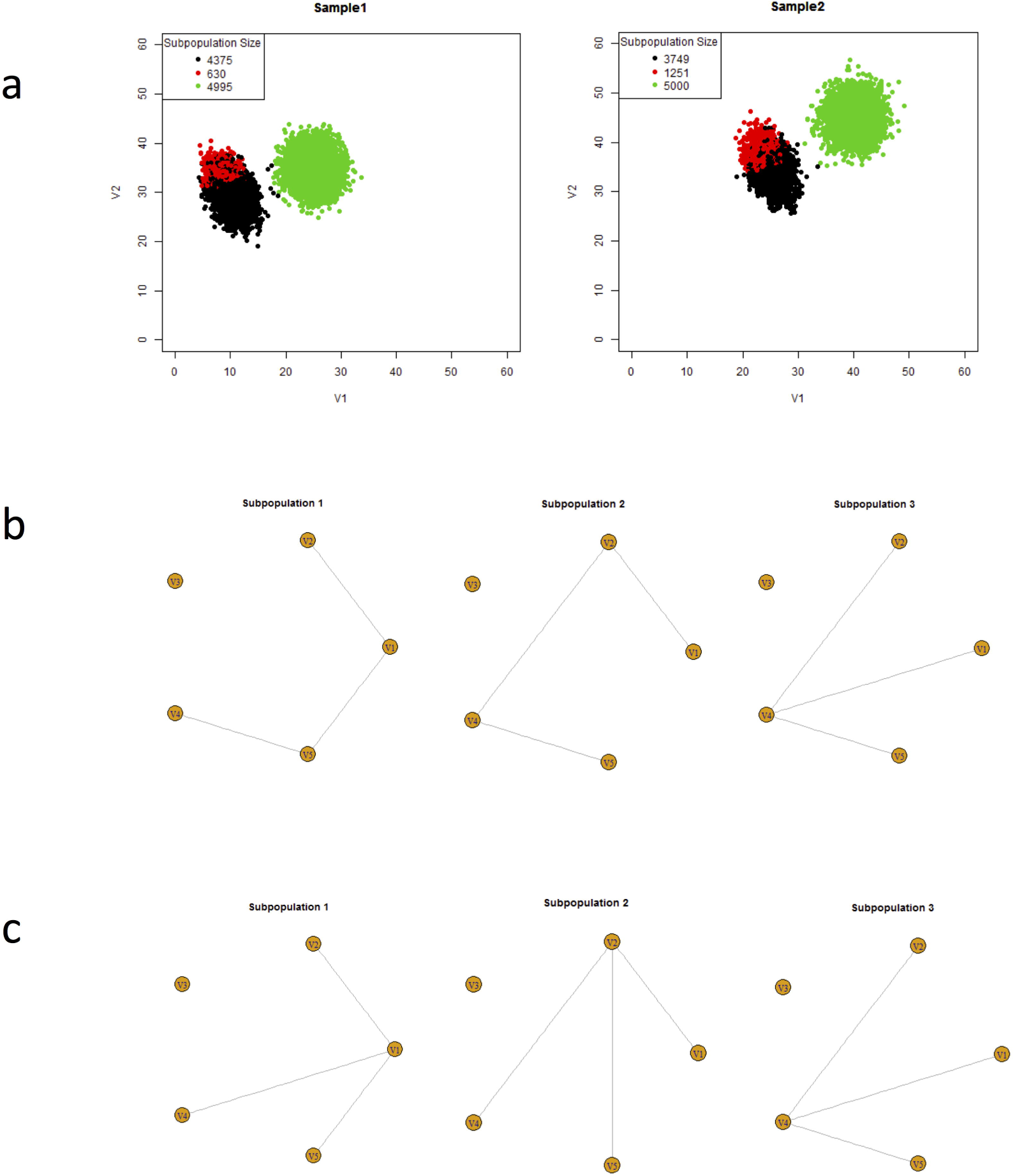
Calculation of sample clusters and their underlying network structures. (a) PAC was used to discover several subpopulations per sample without advanced knowledge of the exact number of subpopulations. (b-c) The networks of the subpopulations in both samples discovered in (a). Networks can be grouped by similarities to organize the subpopulations across samples; the alignment is based on Jaccard dissimilarity network structure characterization matrix; dendrogram of the hierarchical clustering results.

**Fig 8.**
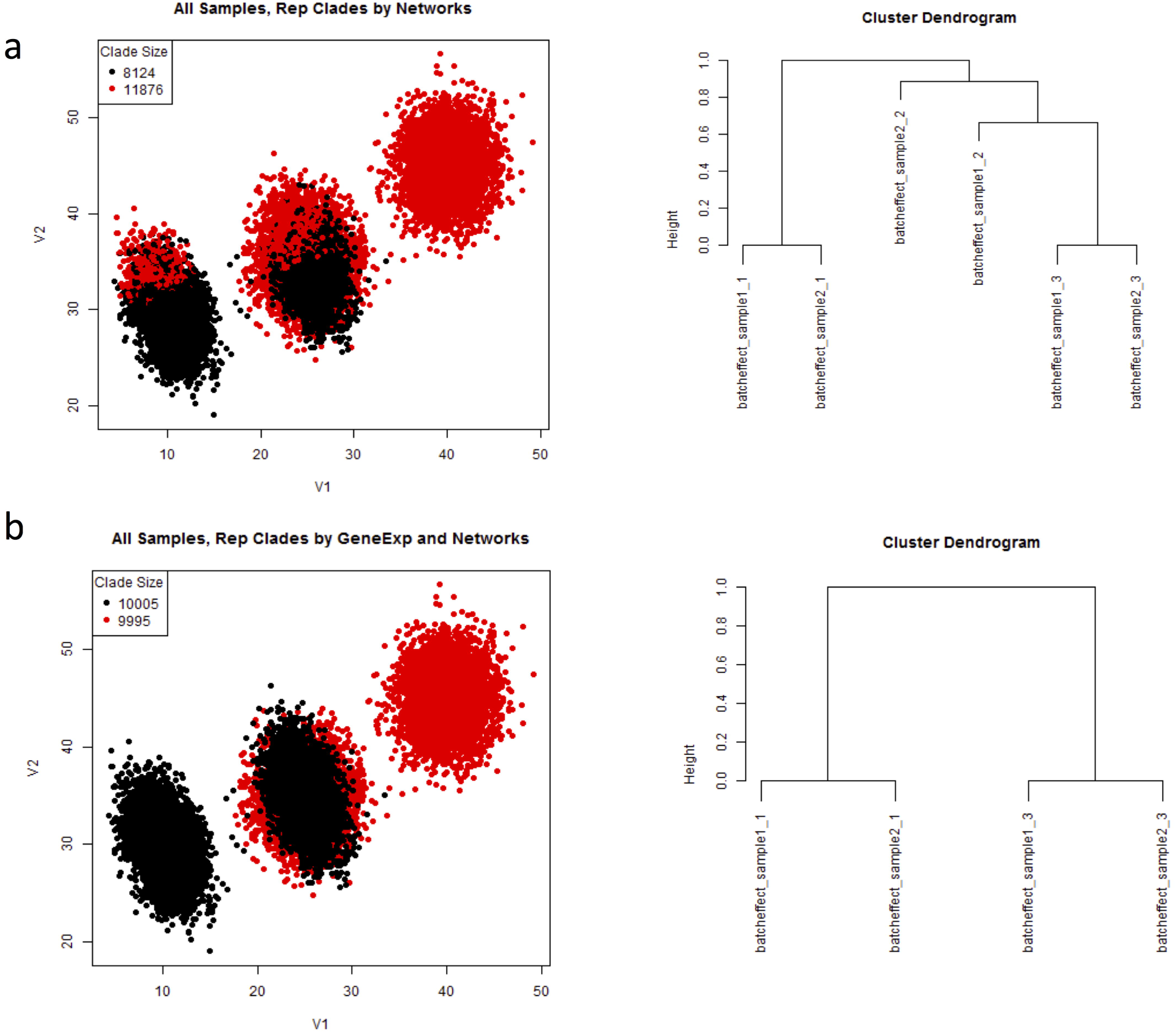
Resolution of batch effects for simple batch effect scenario. (a) Resolution of batch effect by networks of all subpopulations discovered. (b) Resolution of batch effect first by network structures of larger subpopulations and then by merging smaller subpopulations into the aligned clades.

Next we consider the case with dynamic evolution of subpopulations that models the treatment-control and perturbation studies. The interesting information is in tracking how subpopulations change over the course of the experiment. In the simulation, we have generated two subpopulations that nearly converge in mean expression profile over the time course (Fig 9). The researcher could lose the dynamic information if they were to combine the samples for clustering analysis. As in the previous case, we could use PAC to learn several subpopulations per sample (Fig 10). Then, with the assumption that there are two evolving clusters from data exploration, we align the subpopulations to construct clades of same and/or similar subpopulations (Fig 11 left panel) based on the network structural information (S3 Fig). With network and expression level information in the alignment process, the two subpopulations or clades can be resolved naturally (Fig 11 right panel).

**Fig 9.**
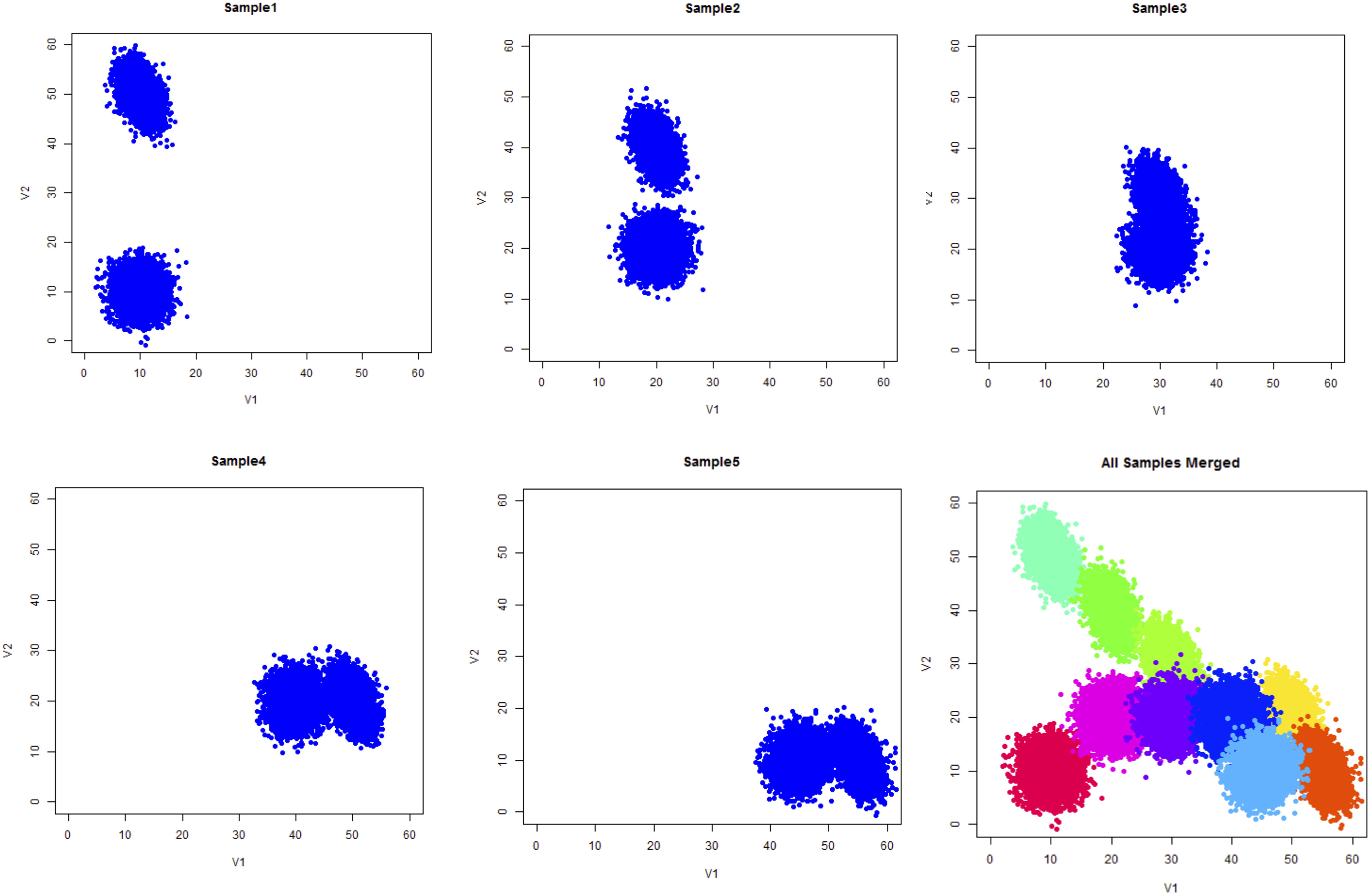
Ground truth of simulated dynamic batch effect samples. Two subpopulations, in blue color, almost converge in time by mean shifts.

**Fig 10.**
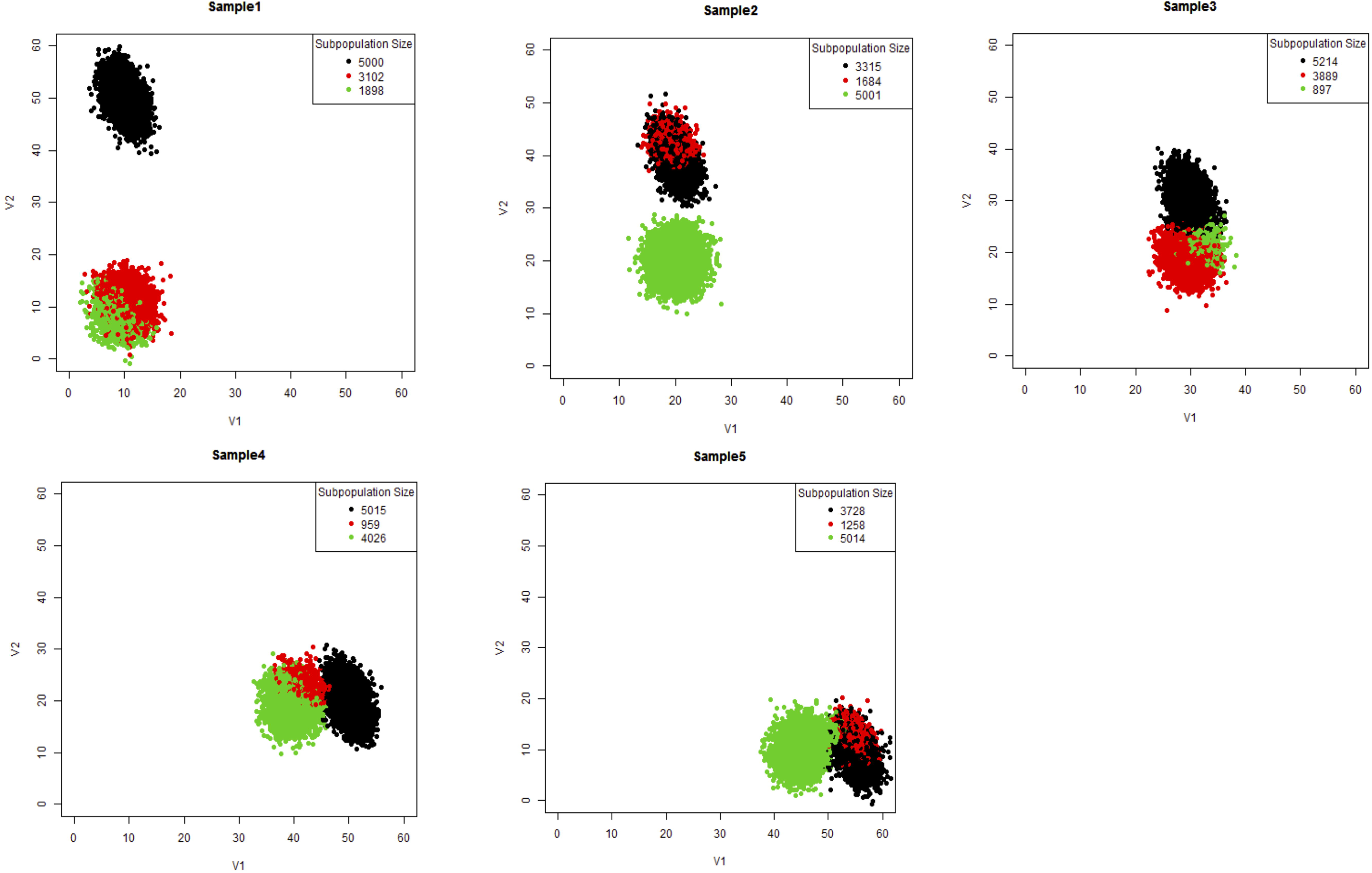
PAC on dynamic batch effect scenario. PAC discovers several subpopulations per sample without advanced knowledge of the number of subpopulations present.

**Fig 11.**
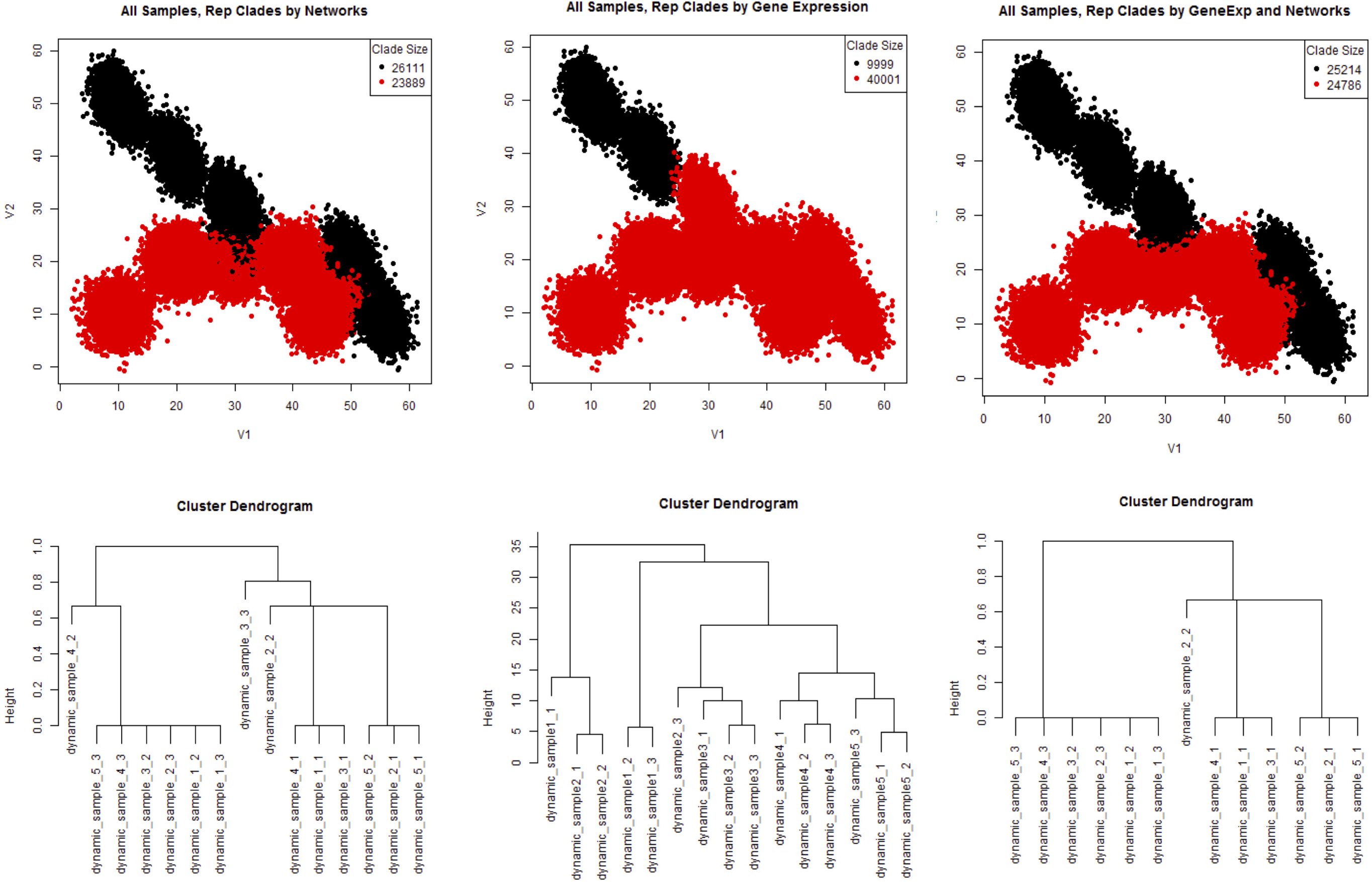
PAC-MAN results for dynamic batch effect scenario. Comparison of PAC-MAN results between representative clades (number of clades set to 2). Using network structures (left panel) or expression information (middle panel) alone does not resolve the dynamic information. On the other hand, the dynamic information is resolved first by alignments of networks of larger subpopulations and then by merging smaller subpopulations into the aligned clades (right panel).

### Network and expression alignment is better than network or expression alignment alone

With networks in hand, we could further characterize the relationships between subpopulations across samples. However, the alignment process needs to work well for true linkage to be established. We could align by network alone, by expression (or marker) means, or both. Figs 8 and 11 present these alternatives in comparison. By using all the subpopulation networks, the results still contain subsets of misplaced cells (Figs 8a and 11 left panel). This is because small clusters of cells have noisy underlying covariance structure; therefore, the networks cannot be accurately inferred. These structural inaccuracies negatively impact the network clustering. The (mean) marker level approach also does not work well (Fig 11 center panel) due to the subpopulation mean shifts across samples. On the other hand, the sequential approach works well (Figs 8b and 11 right panel). In the sequential approach, larger (>1500 in batch effect case; >1000 in dynamic case) subpopulations’ networks are utilized for the initial alignment process. Next, the smaller subpopulations, which have noisy covariance, are merged with the closest larger, aligned subpopulations. Thus, more subpopulations could be discovered upstream (in PAC), and the network alignment would work similarly as the smaller subpopulations, which could be fragments of a distribution, do not impact the alignment process (S4a Fig and S4b Fig). Moreover, in the network inference step, unimportant edges can negatively impact the alignment process (S4c Fig) in the network-alone case. Biologically, this means that edges that do not constrain or define the cellular state should not be utilized in the alignment of cellular states. Effectively, the threshold placed on the number of edges in the network inference controls for the importance of the edges. Thus, the combined alignment approach works well and allows moderate over-saturation of cellular states to be discovered in the PAC step so that no advance knowledge of the exact number of subpopulations is necessary.

### PAC-MAN efficiently outputs meaningful data-level subpopulations for mouse tissue dataset

We use the recently published mouse tissue dataset[11] to illustrate the multi-sample data analysis pipeline. The processed dataset contains a total of more than 13 million cell events in 10 different tissue samples, and 39 markers per event (S2 Table). The original research results centered on subpopulations discovered from hand-gating the bone marrow tissue data to find ‘landmark’ subpopulations; the rest of the data points were clustered to the most similar landmark subpopulations. While this enables the exploration of the overall landscape from the perspective of bone marrow cell types within an acceptable time frame, a significant amount of useful information from the data remains hidden; a larger dataset would make it infeasible to analyze by manual gating and existing computational tools to learn the relationships of the cellular states among all samples. In addition, a natural question is how well do the bone marrow cell types represent the whole immune system?

In contrast to the one-sample perspective, using d-PAC-MAN, the fastest approach by our comparison results, we can perform subpopulation discovery for each sample automatically and then align the subpopulations across samples to establish dataset-level cellular states. On a standard Core i7-44880 3.40GHz PC computer, the single-thread data analysis process with all data points takes about one hour to complete, which is much faster than alternative methods. With multi-threading and parallel processing, the data analysis procedure can be completed very quickly. As mentioned earlier, PAC results for the bone marrow subsetted data from this dataset matches closely to that of the hand-gated results. This accuracy provides confidence for applying PAC to the rest of the dataset.

Figs 11-12 show the t-sne plots for subpopulation discovered (top panel of each sample) and the representative subpopulation established (bottom panel of each sample) for the entire dataset. In the PAC discovery step, we learn 35 subpopulations per sample without advance knowledge of how many subpopulations are present. This moderate over-partitioning of the data samples leads to a moderate heterogeneity in the t-sne plots. Next, the networks are inferred for the larger subpopulations (with number of cell events greater than 1000), and the networks are aligned for all the tissue samples. We output 80 representative subpopulations or clades for the entire dataset to account for the traditional immunological cellular states and sample-specific cellular states present. Within samples, the subpopulations that cluster together by network structure are aggregated. The smaller subpopulations (not involved in network alignment) are either merged to the closest larger subpopulation or establish their own sample-specific subpopulation by expression alignment; small subpopulations were clamped with larger clades by grouping the subpopulations into 5 clusters per sample based on the means (of marker signal). The representative subpopulations (90 total) follow the approximate distribution of the cell events on the t-sne plots and the aggregating effect cleans up the heterogeneities due to over-partitioning in the PAC step.

**Fig 12.**
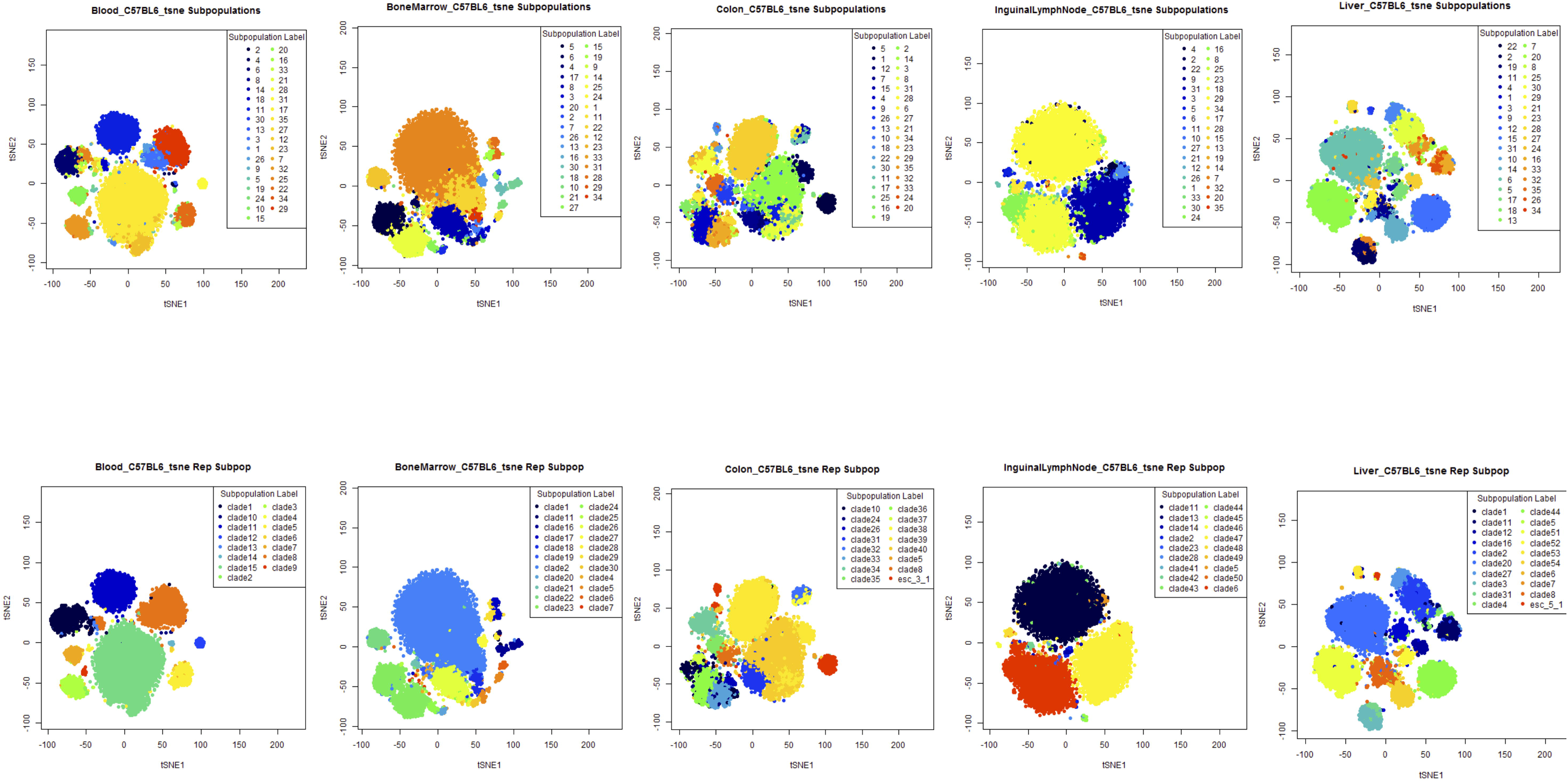
Visualization of PAC-MAN results for Blood, Bone Marrow, Colon, Inguinal Lymph Node, and Liver samples. Each t-sne plot was generated using 10,000 randomly drawn cell events from each mouse tissue sample. The results from PAC (top panel) and MAN (bottom panel) steps are presented as a pair. Initial PAC discovery was set to 35 subpopulations without advanced knowledge of the number of subpopulations in each sample. In MAN, 80 network clades were outputted, and the cellular states are defined by expression (marker signal), network structure, and dataset-level variation. This composite definition naturally aggregates the initial 35 subpopulations to yield smaller number of subpopulations in less variable samples.

**Fig 13.**
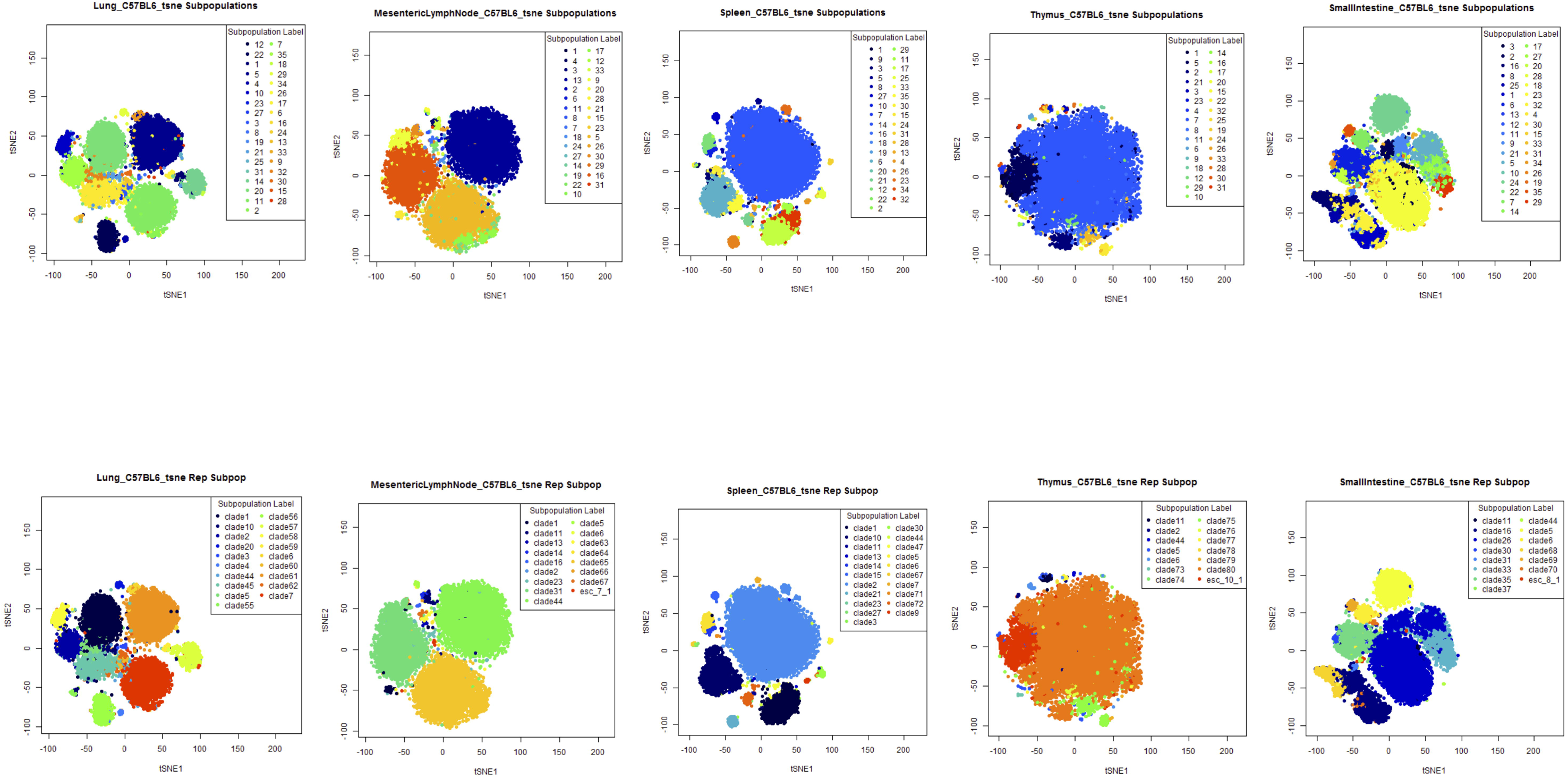
Visualization of PAC-MAN results for Lung, Mesenteric Lymph Node, Spleen, Thymus, and Small Intestine samples. The settings and descriptions are the same as those in Fig 12. Continuation of visualization of PAC-MAN results for the mouse tissue data.

The cell type clades are the representative subpopulations for the entire dataset, and they could either be present across samples or in one sample alone. Their distribution is visualized by a heatmap (Fig 14). While the bone marrow sample contains many cell types, only a subset of them are directly aligned to cell types in other samples, which means using the bone marrow data as the reference point leaves much information unlocked in the dataset. Therefore, the data suggests that the bone marrow cell types are not adequate in representing all cell types in the immune system. The cell types in the blood and spleen samples have more alignments with cell types in other samples. The lymph node samples share many clades; the small intestine and colon samples also share many clades, probably due to closeness in biological function. The thymus sample has few clades shared with other samples, which may be due to its functional specificity.

**Fig 14:**
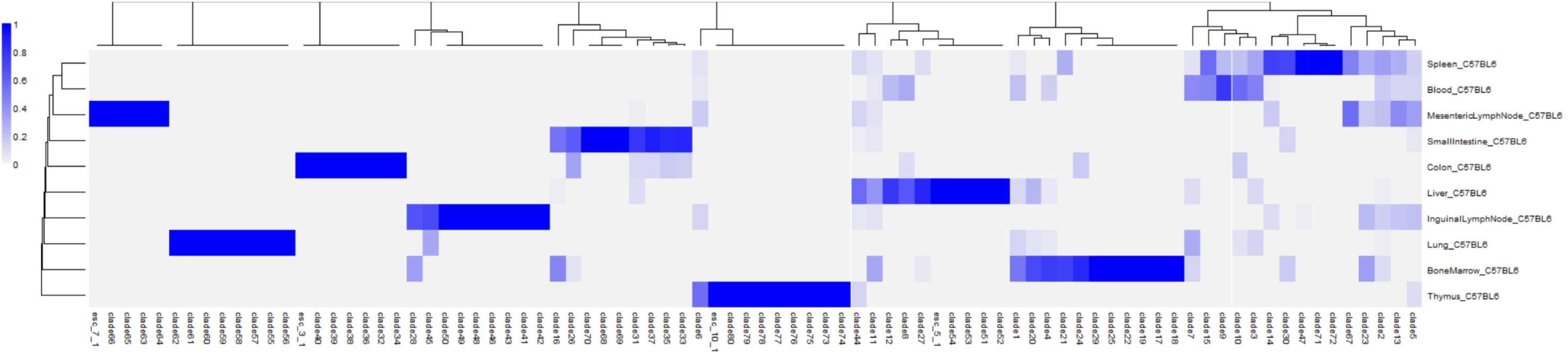
Heatmap of clade proportions across the tissue samples. Sample-specific clades have a value of 1, while shared clades have proportions spread across different samples. Physiologically similar samples share more clades.

PAC-MAN style analysis can be applied to align the tissue subpopulations by their means instead of network similarities (S5 Fig). As done previously, representative clades (88 total) were outputted. The same aggregating effect is observed (S5a Fig), and this is due to the organization from dataset-level variation in the means. Comparing to the network alignment, the means linkage approach has slightly more subpopulations per sample; the subpopulation proportion heatmap (S5b Fig) shows more linking. Although the bone marrow sample subpopulations co-occur in the same clades slightly more with other sample subpopulations, this sample does not co-occur with many clades in the dataset. Thus, a PAC-MAN style analysis with means linkage also harvests additional information from the entire dataset.

To compare the network and means approaches with PAC-MAN, we study the F-measure and p-measure results with 88 total clades from each approach. The overall F-measure with all cell events is 0.7969 and the overall F-measure with clades assignments of PAC-discovered subpopulations is 0.3143. The two F-measure values suggest that the assignment of PAC-discovered subpopulations is more consistent for larger subpopulations.

To illustrate the assignment purities, the p-measures are computed for the following two cases. 1) Network clade assignment is the basis (network-justified), similar to the ground truth in the clustering comparisons previously; or 2) means clade assignment is the basis (means-justified) (S4 Table). P-measure cutoff is set at 0.3 (to remove unreliable comparisons) to obtain purer clade assignments. In the network-justified case, PAC subpopulations with more than 0.3 in p-measure constitute 93.44 *%* of all cell events. In the means-justified case, PAC subpopulations with more than 0.3 in p-measure constitute 92.67 % of all cell events. Furthermore, if the p-measure cutoff were to increase to 0.5, the percentages of cells left for the network-justified and mean-justified cases are 6.25% and 75.16%, respectively. The network-justified case yields drastically lower numbers of cell events in the purer PAC subpopulations because the means approach has more heterogeneity in the linkages (defined as PAC-subpopulation participants in each shared clade with size of at least 2). In fact, the network approach has 100 linkages while the means approach has 209 linkages. Therefore, the extra linkages in the means approach would yield greater impurities in the network-justified case. The linkage plot (S6a Fig) shows that the low linkages occur slightly more frequently for the network approach. One consequence is that the network approach aggregates PAC subpopulations within sample more frequently; for instance, in the thymus sample, the network approach yields 14 clades while the means approach yields 21 clades.

After aggregating, the clade sizes (with unique participants per sample) are plotted (S6b Fig). The network approach tends to find fewer linkages, as more clades have sizes of less than 4, while the means approach has more clades than the network approach with clade sizes greater than 4. The network approach is more conservative due to the additional constraints from network structures. Conventionally, in the cytometry field, only the means are considered in the definition of cellular states. Assuming the absence of batch and dynamic effects, the researcher could view the purer shared clade assignments in the network-justified case (general agreement between constrained network approach and means approach) as more reliable candidates of cross-sample relationships to investigate in future experiments (S6c Fig).

Hence, the network alignment approach is in agreement that of the means approach, with network alignment being more stringent in the establishment of linkages. The network PAC-MAN approach defines cellular states with the additional information from network structures, and it has the effect of constraining the number of linkages between samples while finding linkages for subpopulations that are distant in their means.

### Network hubs provide natural annotations

To further characterize the cell types, we annotate the clades within each sample using the top network hub markers, which constrain the cellular states. The full annotation, along with mean average expression profiles, is presented in S3 Table. The clade information is presented in the ClusterID column. The annotations for cells across different samples but within the same clades share hub markers. For example, in clade 1 for the blood and bone marrow samples, the cells share the hub markers Ly6C and CD11b. In the bone marrow sample, one important set of subpopulations is the hematopoietic stem cell subpopulations. One such subpopulation is present as clade 18 with the annotation CD34-CD27-cKit-Sca1 and is about 1.87 percent in the bone marrow sample. Clade 18 is only present in the bone marrow sample, indicating that the PAC-MAN pipeline defines this as a sample-specific and coherent subpopulation using dataset-level variation. The thymus contains a large subpopulation (84.07 percent) that is characterized as CD5-CD4-CD43-CD3, suggesting it to be the maturing T-cell subpopulation.

## Conclusion

We have presented the PAC-MAN data analysis pipeline. This pipeline was designed to remove major roadblocks in the utilization of existing and future CyTOF datasets. First, we established a quick and accurate clustering method that closely matches expert gating results; second, we demonstrated the management of multiple samples by handling mean shifts and batch effects across samples. The alignment allows researchers to find relationships between cells across samples without resorting to pooling of all data points. PAC-MAN allows the cytometry field to harvest information from the increasing amount of CyTOF data available. It is important to standardize multi-sample data analysis with automation so that discoveries based on multi-sample CyTOF datasets from different laboratories do not depend on the experts’ manual gating strategies and the grouping of subpopulations that is constrained by non-systematic computations. Furthermore, due to PAC-MAN’s generality, this pipeline can be utilized to analyze large datasets of high-dimension beyond the cytometry field.

## Materials and Methods

### Partition-assisted clustering has two parts

1) Partitioning: a partition method (BSP[5] or DSP[7]) is used to learn N initial cluster centers from the original data.

2) Post-processing: A small number (m) of k-mean iterations is applied to the rectangle-based clusters from the partitioning, where m is a user-specified number. We used m=50 in our examples. After this k-means refinement, we merge the N clusters hierarchically until the desired number of clusters (this number is user-specified) is reached. The merging is based on a given distance metric for clusters. In the current implementation, we use the same distant metric as in flowMeans[1]. That is, for two clusters X and Y, their distance D(X, Y) is defined as:

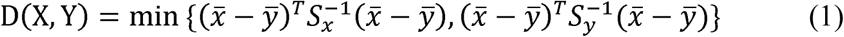

where 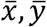 are the sample mean of cluster X and Y, respectively. 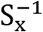 is the inverse of the sample covariance matrix of cluster X. 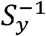 is defined similarly. In each step of the merging process, the two clusters having the smallest pairwise distance will be merged together into one cluster.

### Partition Methods

There are two partition methods implemented in the comparison study: d-PAC and b-PAC. The results are similar, with d-PAC being the faster algorithm. Fig 1a illustrates this recursive process.

d-PAC is based on the discrepancy density estimation (DSP)[7]. Discrepancy, which is widely used in the analysis of Quasi-Monte Carlo methods, is a metric for the uniformity of points within a rectangle. DSP partitions the density space recursively until the uniformity of points within each rectangle is higher than some pre-specified threshold. The dimension and the cut point of each partition are chosen to approximately maximize the gap in uniformity of two adjacent rectangles.

BSP + LL is an approximation inference algorithm for Bayesian sequential partitioning density estimation (BSP)[5]. It borrows ideas from Limited-Look-ahead Optional Pólya Tree (LL-OPT), an approximate inference algorithm for Optional Pólya Tree[8]. The original inference algorithm for BSP looks at one level ahead (i.e. looking at the possible cut points one level deeper) when computing the sampling probability for the next partition. It then uses resampling to prune away bad samples. Instead of looking at one level ahead, BSP + LL looks at h levels ahead (h > 1) when computing the sampling probabilities for the next partition and does not do resampling (Fig 1b). In other words, it compensates the loss from not performing resampling with more accurate sampling probabilities. For simplicity, ‘BSP + LL’ is shortened to ‘BSP’ in the rest of the article.

### F-measure

We use the F-measure for comparison of clustering results to ground truth (known in simulated data, or provided by hand-gating in real data). This measure is computed by regarding a clustering result as a series of decisions, one for each pair of data points. A true positive decision assigns two points that are in the same class (i.e. same class according to ground truth) to the same cluster, while a true negative decision assigns two points in different classes to different clusters. The F-measure is defined as the harmonic mean of the precision and recall. Precision P and recall R are defined as:

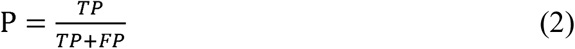

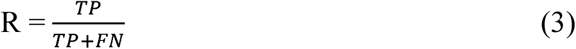
where TP is the total number of true positives, FP is the total number of false positives and FN is the total number of false negatives.

F-measure ranges from 0 to 1. The higher the measure, the more similar the estimated cluster result is to the ground truth. This definition of F-measure is different than that of FlowCAP challenge[2]. The use of co-assignment of labels in this definition is a more accurate way to compute the true positives and negatives.

### Purity-measure (p-measure)

Most of the existing measurements for clustering accuracy aim at measuring the overall accuracy of the entire datasets, i.e. comparing with the ground truth over all clusters. However, we are also interested in analyzing how well a clustering result matches the ground truth within a certain class. Specifically, consider a dataset D with K classes: {C_1_, C_2_, …, C_K_} and a given ground truth cluster labels g, we construct an index called the purity measure, or p-measure for short, to measure how well our clustering result matches g for each class C_i_. This index is computed as follows:

1. For each class C_k_, look for the cluster that has the maximum number of overlapping points with this class, denoted by L_i_k__.
2. Define
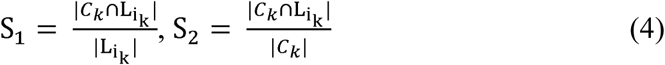
where | · | denotes the number of points in a set.
3. The final P-index for class C_k_ is given by

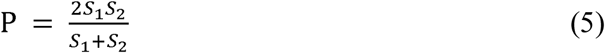

If we were to match a big cluster with a small class, even though the overlapping may be large, S_1_ would still be low since we have divided the score by the size of the cluster in S_1_. In addition, we are interested in knowing how many points in C_k_ are clustered together by L_i_k__, which is measured by S_2_.

### Network construction and comparison

After PAC, the discovered subpopulations typically have enough cells for the estimation of mutual information. This enables the construction of networks as the basis for cell type characterization. Computationally, it is not good to directly use the mutual information networks constructed this way to organize the subpopulations downstream. The distance measure used to characterize the networks could potentially give the same score for different network structures.

Thus, it is necessary to threshold the network edges based on the strength of mutual information to filter out the noisy and miscellaneous edges. In this work, these subpopulation-specific networks are constructed using the MRNET network inference algorithm in the Parmigene [13] R package. The algorithm is based on mutual information ranking, and outputs significant edges connecting the markers. The top *d* edges (*d* is set to be 1x the number of markers in all examples) are used to define a network for the subpopulation. This process enables a careful calculation of the distance measure.

For each pair of subpopulation networks, we calculate a network distance, which is defined as follows. If G_1_ and G_2_ are two networks, let S be the set of shared edges and A be union of the of the edges in the two networks, then we define

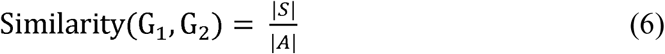
where | · | denotes the size of a set.

This is known as the Jaccard coefficient of the two graphs. The Jaccard distance, or 1-Jaccard coefficient, is then obtained. This is a representation of the dissimilarity between each pair of networks; the Jaccard dissimilarity is the measure used for the downstream hierarchical clustering.

### Cross-sample linkage of subpopulations

We perform agglomerative clustering of the pool of subpopulations from all samples. This clustering procedure greedily links networks that are the closest in Jaccard dissimilarity, and yields a dendrogram describing the distance relationship between all the subpopulations. We cut the dendrogram to obtain the *k* clades of subpopulations. Subpopulations from the same sample and falling into the same clade are then merged into a single subpopulation (Fig 5). This merging step has the effect of consolidating the over-partitioning in the PAC step. No merging is performed for subpopulations from different samples sharing the same clade. In this way, we obtain *k* clades of subpopulations, with each clade containing no more than one subpopulation from each sample. We regard the subpopulations within each clade as being linked across samples.

In the above computation, only subpopulations with enough cells to define a stable covariance are used for network alignment via the Jaccard distance; the rest of the cell events from very small subpopulations are then merged with the closet clade by marker profile via distance of mean marker signals. If the small subpopulations are distant from the defined clades, then a new sample-specific clade is created for these small subpopulations.

### Annotation of Subpopulations

To annotate the cellular states, we first apply PAC-MAN to learn the dataset-level subpopulation/clade labels. Next, these labels are used to learn the representative/clade networks. The top hubs (i.e. the most connected nodes) in these networks are used for annotation. This approach has biological significance in that important markers in a cellular state are often central to the underlying marker network, which is analogous to important genes in gene regulatory networks; these important markers have many connections with other markers. If the connections were broken, the cell would be perturbed and potentially driven to other states.

### Running Published Methods

To run t-SNE [14] a dimensionality reduction visualization tool, we utilized the scripts published here (https://lvdmaaten.github.io/tsne/). Default settings were used.

To run SPADE, we first converted the simulated data to fcs format using Broad Institute’s free CSVtoFCS online tool in GenePattern[15] (http://www.broadinstitute.org/cancer/software/genepattern#).

Next, we carried out the tests using the SPADE package in Bioconductor R[16] (https://bioconductor.org/packages/release/bioc/html/spade.html.

To run flowMeans, we carried out the tests using the flowMeans package in Bioconductor R[1] (https://bioconductor.org/packages/release/bioc/html/flowMeans.html.

In the comparisons, we selected only cases that work for all methods to make the tests as fair as possible.

To calculate the mutual information of the subpopulations, we use the infotheo R package (https://cran.r-proiect.org/web/packages/infotheo/index.html.

To run network inference, we use the mrnet algorithm in the parmigne R package [13]. (https://cran.r-proiect.org/web/packages/parmigene/index.html.

### Code Availability

The PAC R package can be accessed at: https://cran.r-proiect.org/web/packages/PAC/index.html

### Simulated Data for Clustering Analysis

To compare the clustering methods, we generated simulated data from Gaussian Mixture Model varying dimension, the number of mixture components, mean, and covariance. The dimensions range from 5 to 39. The number of mixture components is varied along each dimension. The mean of each component was generated uniformly from a d-dimensional hypercube; we generated datasets using hypercube of different sizes, but kept all the other attributes the same. The covariance matrices were generated as *AA^T^*, where *A* is a random matrix whose elements were independently drawn from the standard normal distribution. The sizes of the simulated dataset range from 100k to 200k.

The simulated data are provided as (Datasets 1-6). Datasets 1-4 are for the PAC part. Dataset 1 contains data with 5 dimensions; Dataset 2 contains data with 10 dimensions; Datasets 3a and 3b contain data with 20 dimensions; and Datasets 4a and 4b contain data with 35 dimensions. The ground truth labels are included as separate sheets in each dataset.

When applying flowMeans, SPADE, and the PAC to the data, we preset the desired number of subpopulations to that in the data to allow for direct comparisons.

### Gated Flow Cytometry Data

Two data files were downloaded from the FlowCAP challenges[2]. One data file is from the Hematopoietic stem cell transplant (HSCT) data set; it has 9,936 cell events with 6 markers, and human gating found 5 subpopulations. Another data file is from the Normal Donors (ND) data set; it has 60,418 cell events with 12 markers, and human gating found 8 subpopulations. The files are the first (‘001’) of each dataset. These data files were all 1) compensated, meaning that the spectral overlap is accounted for, 2) transformed into linear space, and 3) pre-gated to remove irrelevant events. We used the data files without any further transformation and filtering. When applying flowMeans, SPADE, and the PAC to the data, we preset the desired number of subpopulations to that in the data to allow for direct comparisons.

### Gated Mass Cytometry Data

Human gated mass cytometry data was obtained by gating for the conventional immunology cell types using the mouse bone marrow data recently published[11]. The expert gating strategy is provided as Fig S1. The gated sample subset contains 64,639 cell events with 39 markers and 24 subpopulations and it is provided as Dataset 7.

To test the performance of different analysis methods, the data was first transformed using the asinh(x/5) function, which is the transformation used prior to hand-gating analysis; For SPADE analysis, we utilize the asinh(x/5) option in the SPADE commands. The post-clustering results from flowMeans, SPADE, b-PAC, and d-PAC were then subsetted using the indexes of gated cell events. These subsetted results are compared to the hand-gated results.

### Simulated Data for MAN Analysis

To test the linking of subpopulations, we generated simulated data from multivariate Gaussian with preset signal levels and randomly generated positive definite covariance matrices. There are two cases, batch effect and dynamic. Each simulated sample file has five dimensions, with two of these varying in levels; these are the dimensions that are visualized. Dataset 5 contains the data for general batch effects case and Dataset 6 contains the data for dynamic effects case. The ground truth labels are included as separate sheets in each dataset.

#### General batch scenario

Sample 1 represents data from an old instrument (instrument 1) while sample 2 represents data from a new instrument (instrument 2). There are two subpopulations per sample. These two subpopulations are the same, but their mean marker levels shifted higher up in sample 2 due to higher sensitivity of instrument 2 (Fig 6a). The subpopulations have different underlying relationships between the markers. In this simulated experiment, five markers were measured. Out of the five markers, two markers show significant shift, and we focus on these two dimensions by 2-dimensional scatterplots. In Fig 6a, the left subpopulation in sample 1 is the same as the left subpopulation in sample 2; the same with the right subpopulation. The same subpopulations were generated from multivariate Gaussian distributions with changing means with fixed covariance structure.

#### Dynamic scenario

Dynamic scenario models the treatment-control and perturbation studies. In the simulation, we have generated two subpopulations that nearly converge over the time course (Fig 9). The researcher could lose the dynamic information if they were to combine the samples for clustering analysis. The related subpopulations were generated from multivariate Gaussian distributions with changing means with fixed covariance structure.

### Raw CyTOF Data Processing

The researcher preprocesses the data to 1) normalize the values to normalization bead signals, 2) de-barcode the samples if multiple barcoded samples were stained and ran together, and 3) pre-gate to remove irrelevant cells and debris to clean up the data[10,17]. Gene expressions look like log-normal distributions[18]; given the lognormal nature of the values, the hyperbolic arcsine transform is applied to the data matrix to bring the measured marker levels (estimation of expression values) close to normality, while preserving all data points. Often, researchers use the asinh(x/5) transformation, and we use the same transformation for the CyTOF datasets analyzed in this study.

### Mouse Tissue Data

In the Spitzer et al., 2015 dataset[11], three mouse strains were grown, and cells were collected from different tissues: thymus, spleen, small intestine, mesenteric lymph node, lung, liver, inguinal lymph node, colon, bone marrow, and blood. In each experiment, 39 expression markers were monitored. The authors used the C57BL6 mouse strain as the reference[11]; the data was downloaded from Cytobank, and we performed our analysis on the reference strain.

First, all individual samples were filtered by taking the top 95% of cells based on DNA content and then the top 95% of cells based on cisplatin: DNA content allows the extraction of good-quality cells and cisplatin level (low) allows the extraction of live cells. Overall, the top 90% of cell events were extracted. The filtered samples were then transformed by the hyperbolic arcsine (x/5) function, and merged as a single file, which contains 13,236,927 cell events and 39 markers per event (S2 Table).

Using PAC-MAN, we obtained 35 subpopulations in each sample then 80 clades for the entire dataset. The 80 clades account for the traditional immune subpopulations and sample-specific subpopulations. Small subpopulations not used in alignment are later merged into the closest clades; this is done by performing hierarchical clustering with the marker signals to obtain 5 “expression” subclades per sample. Subsequently, any clade with less than 100 cells is discarded. Subpopulation proportion heatmap was plotted to visualize the subpopulation-specificities and relationships across the samples. Finally, annotation was performed using the hub markers of each representative subpopulation in each sample.

## Acknowledgements

We thank the members of Wong Lab, in particular Tung-yu Wu, Chen-yu Tseng and Kun Yang, for critical feedback.

## Supporting Information

**S1 Fig. Gating strategy of CyTOF data for methods comparison.** Biaxial gating hierarchy for the mouse bone marrow CyTOF dataset. Gating strategy that was used to find 24 reference populations in the mouse bone marrow CyTOF data. Pre-gating step involved removal of doublets, dead cells, erythrocytes and neutrophils. Non-neutrophils population was either subject to cluster analysis by computational tools or subsequent gating. Dotted boxes represent 24 terminal gates that were selected as reference populations for the comparison analysis.

**S2 Fig. Subpopulation purity of simulated and real CyTOF data.** (a) Subpopulation-specific purity plot of 35-dimensional simulated data with 10 subpopulations. The blue points denote the differences between the p-measures of the partition-based method (either d-PAC or b-PAC) and flowMeans, while the red points denote the p-measure differences between the partition methods and SPADE. The horizontal line at 0 means no difference between the methods. Most of the blue and red points are above 0, indicating that the PAC generates purer subpopulations compared to the ground truth. The two subplots are very similar, which means that d-PAC and b-PAC give very similar p-measures. More precisely, the sum of differences between d-PAC and flowMeans and d-PAC and SPADE are 0.85 and 1.09, respectively; and the overall difference between b-PAC and flowMeans and b-PAC and SPADE are 0.84 and 1.08, respectively.

(b) Subpopulation-specific purity plot of the hand-gated CyTOF data. The same convention is used as in (S2a Fig). Again, more blue and red points are above 0, indicating that the partition-based methods generate purer subpopulations compared to the ground truth. There is a cluster of points below 0 occurring in the middle of the plot, suggesting that flowMeans and SPADE capture the mid-size subpopulations more similar to hand-gating than the partition-based methods. More specifically, flowMeans does better (p-measure difference of 0.1 or better; difference of less 0.1 is considered practically no difference) with finding subpopulations of GMP, CD8 T cells, MEP, CD4 T cells (compared to d-PAC), and Plasma cells, while SPADE does better with CD19+IgM-B cells, NK cells (compared to d-PAC), CD8 T cells, NKT cells, Basophils, Short-Term HSC, and Plasma cells. However, overall, PAC has a much better performance, as the absolute sum of points above 0 is higher than that of points below 0. More precisely, the sum of differences between d-PAC and flowMeans and d-PAC and SPADE are 1.21 and 1.45, respectively; and the overall difference between b-PAC and flowMeans and b-PAC and SPADE are 2.06 and 2.31, respectively. The difference table is provided in S1 Table.

**S3 Fig. Networks inferred from subpopulations in the dynamic example simulated dataset.** Fig 9 introduced the dynamic example in which five samples each having 2 true subpopulations captures the almost-convergence of means. Here the underlying network structures for the PAC discovered subpopulations (three per sample) are presented.

**S4 Fig. Comparison between aligning cross-sample subpopulations by network, expression profile, or both.** (a) PAC can be used to discover more subpopulations, with the effect of more partitions from the true clusters. (b) When over-partitioning is present, network or expression profile alone cannot resolve the dynamic (or batch) effects due to noisy covariance for small fragments of distributions. However, first aligning the larger subpopulations with more stable covariance, and thus network structures, and then merge in the smaller subpopulations by expression profile resolves the effects. (c) If more irrelevant edges were introduced, network alignment would fail due to the negative impact of the miscellaneous edges; however, eliminating small subpopulations from the alignment step alleviates the increased edge count problem.

**S5 Fig. PAC-MAN style linkage by means.** (a) t-sne plots of mouse tissue samples colored by representative subpopulations labels from linkage by means. (b) Subpopulation proportion heatmap of clades of samples from linkage by means.

**S6 Fig. Comparison between network and means PAC-MAN.** (a) PAC-discovered subpopulations are aggregated by MAN into clades; the number of PAC subpopulations/clades for the network and means PAC-MAN approaches are plotted. (b) After aggregating shared clades within samples, the number of shared clades for the entire dataset is plotted for the two PAC-MAN approaches. c) Using the network approach results as basis, the clades with strong agreement (high p-measures) with the means PAC-MAN approach are given. The shared clades (present in more than one sample) are reliable candidates for future experiment to find cross-sample relationships.

**S1 Table. Purity (p) Measure Differences in CyTOF Comparison.** p-measure differences in gated CyTOF data analysis comparison. The differences are shown for all the annotated cell subpopulations, which are ordered by their sizes. Overall, the PAC methods give more positive p-measures.

**S2 Table. Sample Sizes in Mouse Tissue CyTOF Dataset.** The numbers of cells in the samples of Spitzer et al., 2015 CyTOF dataset. The data is from the C57BL6 mouse strain and a total of ten tissue samples are present. The raw column shows the number of cells prior to filtering by DNA and cisplatin values. The final cell counts are shown in the filtered file (3rd) column.

**S3 Table. PAC-MAN Subpopulation Characterization Output for Mouse Tissue CyTOF Dataset.** The full set of annotated results, along with mean expressions, subpopulation proportion and counts, are reported.

**S4 Table. Network-justified and means-justified p-measures for Alignments of PAC-discovered Subpopulations.** The PAC-discovered subpopulations were mapped as clades in both the network and means PAC-MAN approaches. The p-measures were calculated for the cases 1) network approach mapping as the basis and 2) means approach mapping as the basis. The comparison is the same in principle to the comparison of labels for clustering methods. The results are ordered by p-measures.

